# Vagal involvement in non-dipping phenotype of Hsd11b2 knockout rats

**DOI:** 10.1101/2022.02.11.480066

**Authors:** Linda J Mullins, Yolanda G S Koutraki, Matthew A Bailey, John J Mullins

## Abstract

The Syndrome of Apparent Mineralocorticoid Excess (SAME) is a hypertensive disorder caused by deficiency of 11b-hydroxysteroid dehydrogenase type 2. Blood pressure is directly influenced by dietary salt intake, but the causes of salt-sensitivity are not fully resolved. We modelled SAME in Fischer 344 rats, using zinc finger nuclease targeting of the Hsd11b2 gene. The F344 genetic background showed modest salt sensitivity: blood pressure increased by ∼6mmHg when diet was switched from control (0.3% Na) to high salt (3% Na) diet. Homozygous knockout (Hsd2^-/-^) rats exhibited severe hypertension on control diet (mean arterial blood pressure of ∼180 mmHg compared to ∼115 mmHg in wild-types) and displayed no dipping in blood pressure in the inactive/sleep phase. They also displayed reduced heart rate (339 bpm versus 384 bpm in F344 controls). Low salt diet (0.03% Na) caused a dramatic fall in Hsd2^-/-^ blood pressure (to ∼141mmHg), restoration of robust circadian variation in blood pressure, and an increase in heart rate (to 364bpm). This was mirrored by a restoration of circadian variation in the Poincare plot descriptor, SD1, suggesting involvement of parasympathetic dysfunction in the non-dipping phenotype. Alpha adrenoceptor blockade with prazosin treatment resulted in a further decrease in blood pressure (to ∼124mmHg), which blunted circadian rhythm, together with an increase in heart rate (to ∼394bpm). This rat model of human hypertension reveals clear links between dietary salt, autonomic nervous system dysfunction, and the non-dipping blood pressure phenotype.

## Introduction

High blood pressure is the major modifiable risk factor for cardiovascular disease. Moreover, salt-sensitivity of blood pressure and a loss of the normal circadian variation (non-dipping) will independently increase cardiovascular risk even in the absence of hypertension. Contributory mechanisms are not fully defined but examination of the rare monogenic hypertensive disorders have revealed that proteins involved with sodium balance are key to the maintenance of BP homeostasis (1-3). The syndrome of apparent mineralocorticoid excess (SAME; OMIM #218030), for example, is caused by a deficiency in the enzyme 11β-hydroxysteroid dehydrogenase type 2 (Hsd11b2), which converts active cortisol (corticosterone in rodents) to inactive cortisone (11-dehydrocorticosterone) (4) in Hsd11b2-expressing tissues such as kidney, colon and placenta. Ligand specificity of the mineralocorticoid receptor (MR) for aldosterone is thus achieved in these tissues. In the absence of Hsd11b2, however, MR is activated illicitly by the highly abundant glucocorticoids, promoting inappropriate sodium re-absorption and potassium secretion in the distal tubule of the nephron. SAME patients are therefore hypertensive and hypokalemic and although the renin-angiotensin-aldosterone system is suppressed, hypertension is strongly salt-sensitive. SAME patients present with short stature, failure to thrive, polydipsia, polyuria and other renal pathologies (5).

Many individuals with salt sensitive hypertension, metabolic syndrome (6) or left ventricular hypertrophy (7) have a high incidence of target organ damage and display disrupted circadian blood pressure rhythms, such as nocturnal non-dipping (where BP falls by less than 10% during the sleep phase) on a high salt diet. Meta-analysis suggests that modest salt reduction reduces blood pressure (8) and this can also rectify non-dipping, restoring the normal circadian variation (9-11).

We previously generated and extensively characterized a mouse Hsd11b2 knockout model (12-16). In the present study, zinc-finger nuclease (ZFN) technology (17,18) was used to generate an Hsd11b2 knockout rat model of SAME (19). These rats demonstrate many of the symptoms of SAME, including reduced size, polydypsia, polyuria, hypertension, sodium retention and hypokalaemia (19). Both mouse and rat Hsd2^-/-^ knockout animals, exhibit aberrations in circadian blood pressure. In the mouse the non-dipping phenotype was shown to reflect changes in NCC expression due to glucocorticoid (20).

Heart rate variability (HRV) determines fluctuations in time intervals between consecutive heart beats (inter-beat intervals or IBI) as the heart responds to physical, environmental or psychological challenges (21). Metrics including time domain, frequency domain and non-linear parameters can be derived from raw blood pressure data. Time-domain parameters quantify the amount of variability in HRV. Frequency domain parameters consider the distribution of heart rate oscillations into ultra-low-, very low-, low-and high-frequency bands. Non-linear parameters quantify the random or unpredictable nature of the time series, and have been linked to dynamics of the autonomic nervous system (ANS) (21). Here we looked at heart rate variability (21) in the rat model, to determine if changes in sympatho-vagal control of blood pressure account for non-dipping. This rat model extends our capabilities to explore the mechanisms underlying SAME, together with non-dipping of blood pressure.

## Methods

### Animal husbandry

All experimental animal studies were undertaken under UK Home Office license, following review by local ethics committee. Rats were maintained in a 12-h light-dark cycle (on at 07.00 h) under controlled conditions of humidity (50 +/-10%) and temperature (21+/-2 °C) and fed standard rat chow (RM1, containing 0.3% Na with soya protein, SDS Ltd., Whitham, Essex, UK) and water ad libitum unless otherwise stated. Fischer, (F344IcoCrl) rats were supplied by Charles River Laboratories. Hsd11b2 knockout rats were generated by zinc-finger nuclease targeting on the Fischer F344 genetic background, as previously described (19), and several indels were identified at the target site.

### Metabolic study

Male rats (groups of 4 to 5) were housed individually in metabolic cages and after an acclimatization period (four days), with standard rat chow (0.3% Na) and water *ad libidum*, urine samples were collected for 5-7 days to establish a baseline. Diet was then changed to either high salt (3% Na) diet or a low salt diet (0.03% Na) for an additional 7 to 10 days. Food and water consumption were measured throughout, and urine samples collected (data not shown). Prazosin (50mg/l) was administered in drinking water where specified.

### Telemetric analyses

Rat telemetry devices (TA11PA-C40; Data Sciences International, St. Paul, Minnesota, USA), were implanted into male rats (minimum size of 250g; groups of 4-5) with the catheter placed below the bifurcation of the renal arteries, in the abdominal aorta, under anaesthesia (isoflurane). Vetegesic was administered post-surgery. Systolic and diastolic blood pressures, together with mean arterial blood pressure (MABP), heart rate, pulse pressure and activity were monitored, and data was collected at 5 kHz for 5 min, every hour. Animals were allowed to recover from surgery for 7 days before the collection of baseline data and subsequent dietary changes as specified. The 24-hour MABP and the day (light phase; L) and night (dark phase; D) MABP were all derived from the raw data. The LD difference was the difference in means of the dark and light phase respectively. The LD phase ratio was the LD difference expressed as a percentage of the night-time average.

### Heart rate variability

After loading the raw DSI pressure waveform data, NaN values (<1% of the dataset) were stripped and replaced using the ‘interpolate between adjacent points’ option. Waveforms were marked on the cleaned data and Inter-beat intervals (IBI) were extracted and exported, 24 hours at a time (24 files per folder). Data was converted from s to ms using the Artifact converter program and folders were run through Artiifact (22), one at a time using the batch processor, in order to strip artifacts from the data. This generated 3 files (.FIG; .PNG (with raw and converted waveforms) and .txt). The corrected .txt files were opened, one by one using Kubios heart rate variability software version 2.0 (Biosignal Analysis and Medical Imaging Group, Department of Physics, University of Kuopio, Kuopio, Finland) (23). Following artifact correction, time-domain, frequency domain and non-parametic parameters were calculated using Kubios. Data was then exported and analyzed for interaction between each parameter, day-night variation in MABP, and genotype or salt diet as appropriate.

### Statistical analyses

Data are presented as the mean +/-SD unless otherwise stated. Interaction plots were generated using Minitab. Mean values were compared using the Student’s t-test. A P<0.05 was considered statistically significant.

## Results

### Telemetric analyses

ZFN targeting of Hsd11b2 generated multiple indels that brought a TGA stop codon into frame, truncating the translation product coded by the Hsd11b2 gene. The deletion of 123bp removed exon2 and part of intron B, causing truncation of the open reading frame (19). The knockout line carrying the 123bp deletion (Δ123) was used for all further analyses.

Homozygous and heterozygous male rats carrying the targeted allele Δ123 (n=4); 16-20 weeks old) were implanted with telemetry devices, to determine their MABP, compared to F344 control littermates. As can be seen (Fig. 1a and Table 1), on standard rat chow (0.3% Na) blood pressure was ∼6mmHg higher in the heterozygous animals than F344 controls (119.83+/-8.53 versus 115.69+/-8.37; p= 0.12), but this was not statistically significant. When switched to a 3% Na diet, both Hsd2^+/-^ animals and controls showed modest salt sensitivity, with a small (∼6mmHg; p=0.012) but significant increase in blood pressure (126.15+/-9.50 versus 121.96+/-8.60; p=0.08). Heterozygotes exhibited a slightly increased light-dark difference (LD diff) and LD phase ratio compared to controls on both the 0.3% Na diet and the 3% Na diet. Interestingly, neither the controls nor the Hsd2^+/-^ animals showed a change in heart rate (0.3%Na: 384+/-22 versus 384+/-28; 3% Na: 387+/-25 versus 384+/-29; not significant; Fig. 1b) or locomotor activity (Fig.1c) on the different diets.

**Figure 1:**
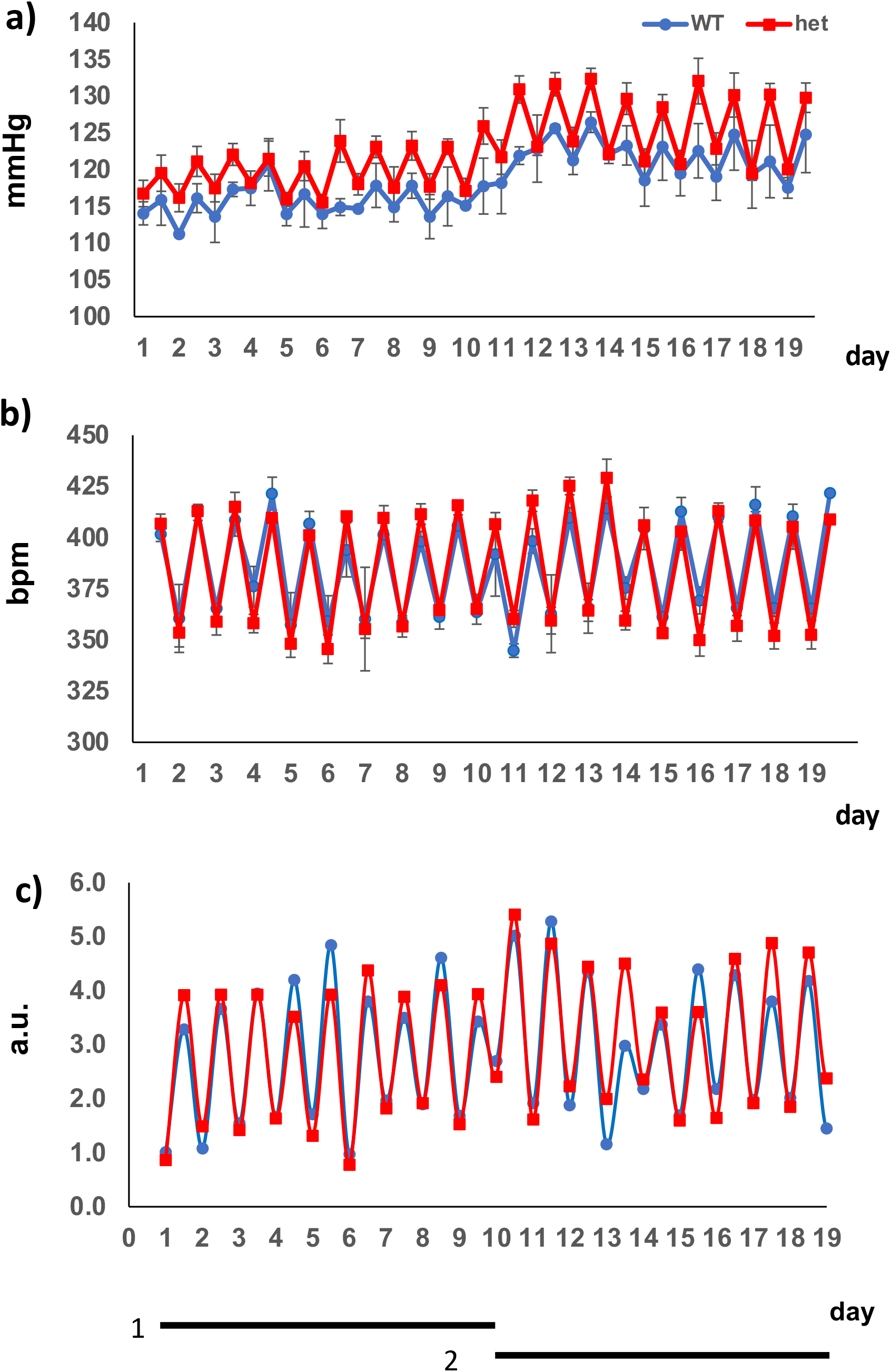
Telemetric analyses a) MABP b) HR and c) locomotor activity analyses of Hsd2^+/-^rat compared to Fischer (F344 WT) controls on (1) 0.3% Na diet and (2) 3% Na diet

**Table 1:** Mean arterial blood pressure and associated light-dark (L/D) variables for F344 wild type and Hsd2^+/-^heterozygotes on normal (0.3%) or high (3%) salt diet

| **Genotype** | **WT (n=4)** | **Hsd2<sup>+/-</sup> (n=4)** |
| --- | --- | --- |
| **Normal salt diet** | **0.3% Na<sup>+</sup>** | **0.3% Na<sup>+</sup>** |
| **24h MABP** | 115.69+/-8.37 | 119.83+/-8.53 |
| **L MABP** | 114.02+/-8.03 | 117.08+/-7.79 |
| **D MABP** | 117.27+/-8.50 | 122.40+/-8.43 |
| **LD difference** | 3.25 | 5.34 |
| **LD phase ratio** | 2.77 | 4.36 |
| **High Salt diet** | **3% Na<sup>+</sup>** | **3% Na<sup>+</sup>** |
| **24h MABP** | 121.96+/-8.60 | 126.15+/-9.50 |
| **L MABP** | 120.03+/-8.00 | 121.75+/-8.07 |
| **D MABP** | 123.68+/-8.81 | 130.50+/-8.62 |
| **LD difference** | 3.65 | 8.75 |
| **LD phase ratio** | 2.95 | 6.7 |

Baseline measurements for homozygous animals (n=4), collected on a standard 0.3% Na diet, revealed that the Hsd2^-/-^ animals were severely hypertensive (MABP of 182.32+/-19.19mmHg compared to control littermates – 115.69+/-8.37mmHg; p=0.003). Moreover, when the day-night variation in MABP was plotted (Fig. 2A – treatment 1; Table 2) it was discovered that the homozygotes did not show any fall in blood pressure during the daytime - characteristic of ‘non-dipping’ in the inactive or sleep period. The diet was changed to a low salt diet (0.03% Na) and MABP immediately dropped by over 40mmHg, over the following 3 days (to 140.82+/-5.84mmHg; p= 1.81e^-26^; Fig. 2a – treatment 2), though not completely down to control blood pressure levels. A robust circadian variation in MABP or dipping was completely restored on the low salt diet (Fig.2a – treatment 2; Table 2).

**Figure 2:**
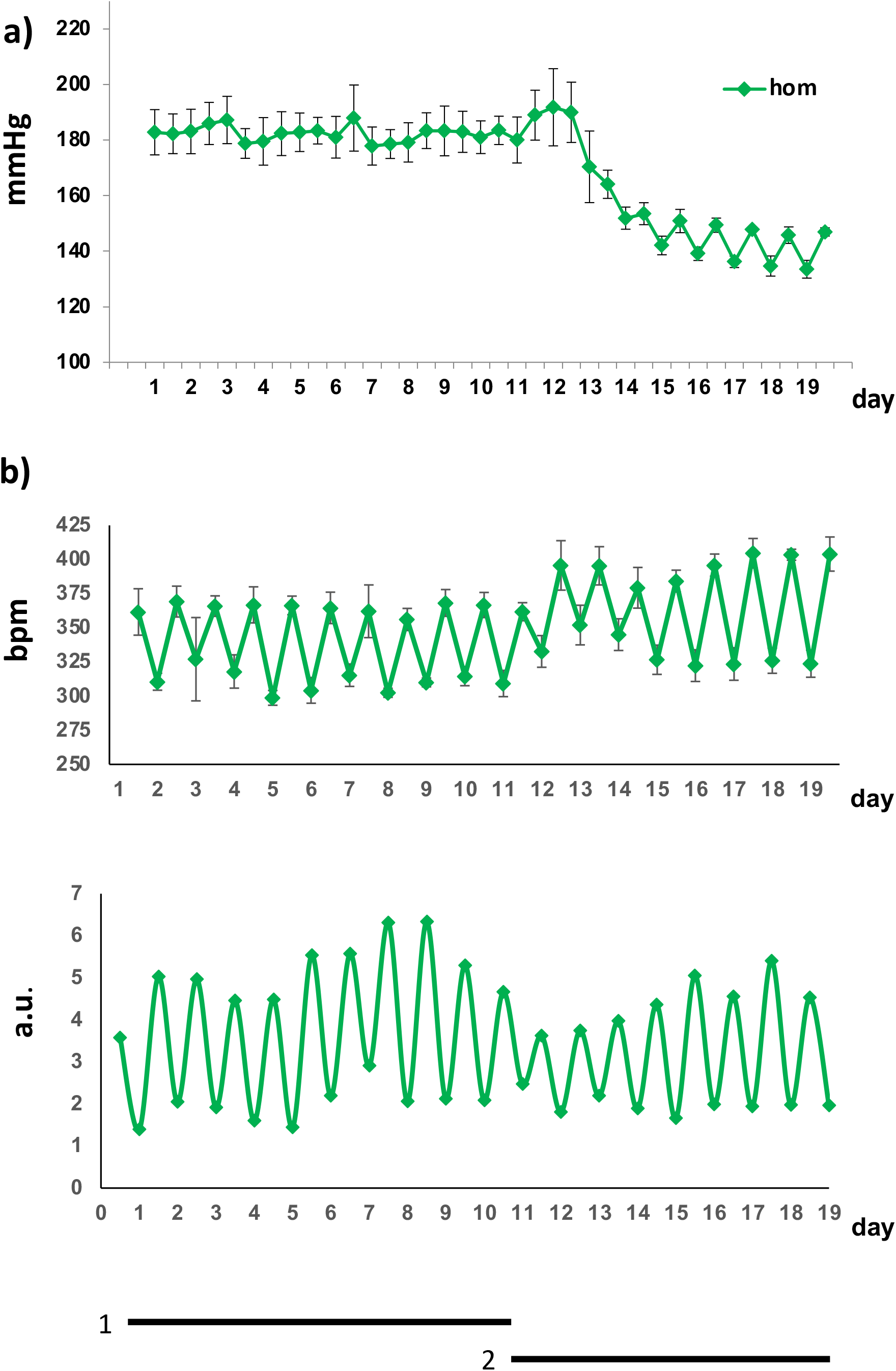
Telemetric analysis showing 12 hourly average of a) MABP, b) HR, and c) locomotor activity of Hsd2^-/-^animals (n=4) on (1) 0.3% Na diet and (2) 0.03% Na diet

**Table 2:** Mean arterial blood pressure and associated light-dark (L/D) variables for F344 wild type and Hsd2^-/-^homozygotes on normal (0.3%) Na diet, compared with Hsd2^-/-^on a 0.03% Na diet. The additional effect of prazosin treatment (PRA; 50mg/l in drinking water) together with 0.03% Na diet on Hsd2^-/-^rats is also shown.

| **Genotype** | **WT (n=4)** | **Hsd2<sup>-/-</sup> (n=4)** | **Hsd2<sup>-/-</sup> (n=4)** | **Hsd2<sup>-/-</sup> (n=4)** |
| --- | --- | --- | --- | --- |
| **diet** | **0.3% Na** | **0.3% Na** | **0.03% Na** | **0.03% Na + PRA** |
| **24h MABP** | 115.69+/-<br>8.37 | 182.32+/-<br>19.19 | 140.82+/-<br>5.84 | 124.23+/-<br>3.25 |
| **L MABP** | 114.02+/-<br>8.03 | 181.97+/-<br>19.67 | 136.54+/-<br>3.01 | 122.30+/-<br>1.34 |
| **D MABP** | 117.27+/-<br>8.50 | 182.56+/-<br>18.88 | 147.11+/-<br>2.37 | 125.52+/-<br>3.08 |
| **LD difference** | 3.25 | 0.59 | 10.57 | 3.22 |
| **LD phase ratio** | 2.95 | 0.32 | 7.19 | 3.56 |

Homozygous Hsd2^-/-^ heart rate (Fig 2b – treatments 1 and 2) showed clear circadian variation on both diets. The change in diet from 0.3% to 0.03% Na, however, led to a significant increase in heart rate (from 339.38+/-28.22 to 363.95+/-34.91 bpm; p=0.024). There was no significant change in locomotor activity (Fig. 2c). Further treatment of Hsd2^-/-^ rats on the 0.03% Na diet with the α1adrenoreceptor blocker, prazosin (50mg/l in drinking water; equates to ∼2-3mg/kg/day; monitored for 5 days) caused an additional increase in heart rate (to 394+/-43 bpm; p=0.03) and a further drop in blood pressure of ∼20mmHg (to 124.23+/-2.04mmHg; p=1.1e^-6^), but the exaggerated dipping phenotype was blunted to control levels (Table 2).

Heart rate variability was compared between control and homozygous animals on 0.3% diet, and between homozygous animals on 0.3% diet and 0.03% diet, to investigate sympatho-vagal involvement. We used Kubios (23) to determine Time domain, Frequency domain and non-linear parameters relating to heart rate variability and looked for any clear interaction between day and night variation of each parameter with salt diet.

Poincare analysis plots each inter-beat interval against the next and gives information about short-term and long-term variability (21,24). The standard deviation of points perpendicular to (SD1) or along the axis (SD2) of the plot distribution relate to vagal (short-term) or sympathetic (long-term) variability respectively. Non-linear parameters calculated by Kubios include SD1 and SD2, approximate entropy and detrended fluctuations. Interaction plots were generated between the day and night averages of each parameter versus salt diet. Only SD1 showed a much greater variation on the low salt (0.03% Na) diet than 0.3% Na diet, reflecting the interactions between the day and night averages of MABP and diet (see Fig. 3a-b).

**Figure 3:**
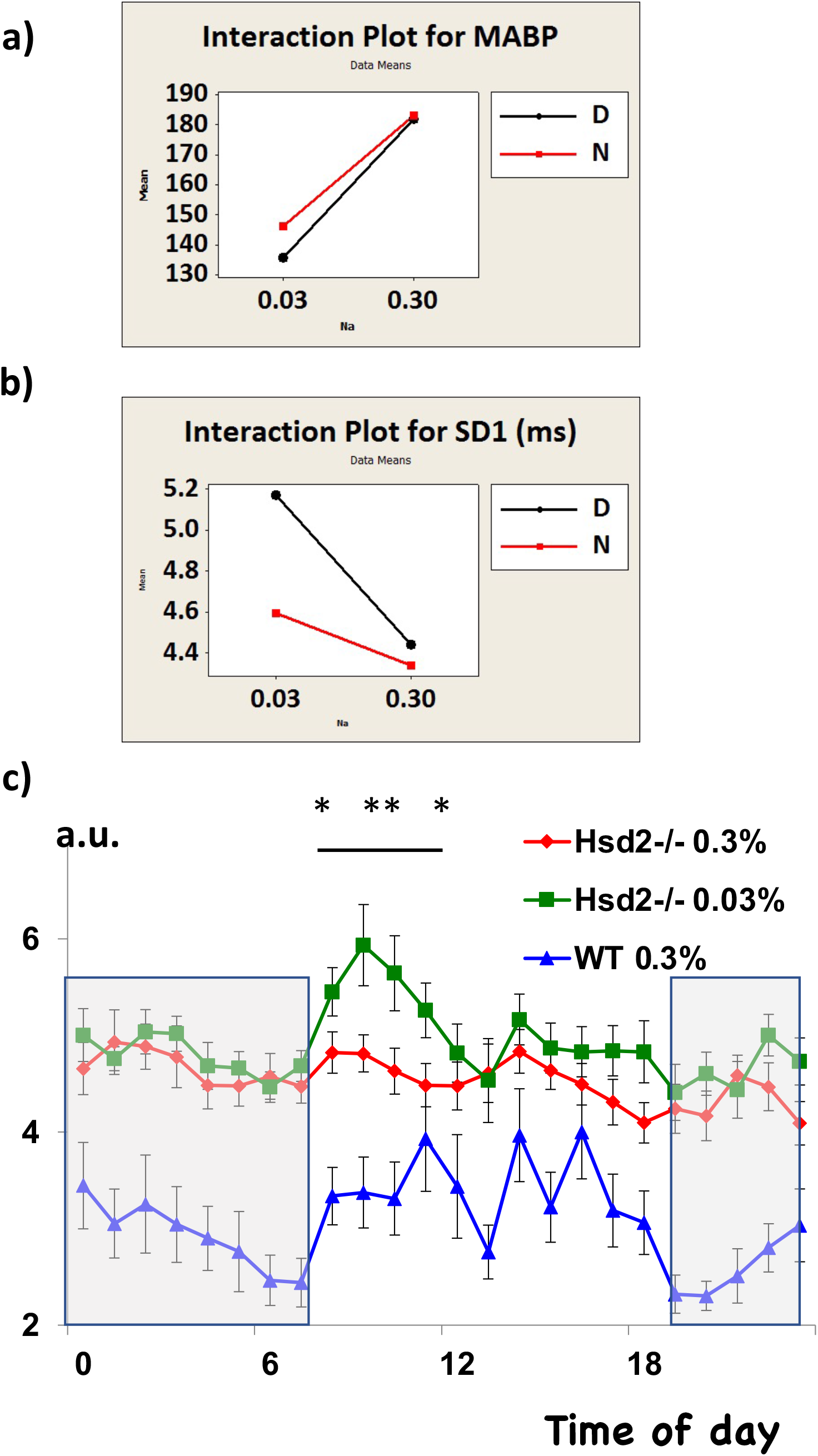
Plots of the day and night averages of Hsd2^-/-^animals showing interactions between a) MABP with Na diet and b) SD1 with Na diet. c) Circadian variation in SD1 of F344 WT and Hsd2^-/-^rats (n=4 per group) on 0.3% Na diet and Hsd2^-/-^on 0.03% Na diet (p *<0.05; **<0.02). Grey area represents dark or active phase

Circadian variation in SD1 was plotted for controls and Hsd2^-/-^ animals on 0.3% diet and compared with Hsd2^-/-^ animals on the 0.03% Na diet (Fig. 3c). Clearly there was minimal circadian variation in SD1 on the 0.3% Na diet, but circadian rhythm of SD1 was restored on the low salt diet, with animals showing a peak in SD1 between 8 am and 11am during the light or inactive period.

## Discussion

We previously used ZFN to generate a rat model of SAME, a human monogenic disorder of salt-sensitive hypertension (19). In this study we examined longitudinal BP traces demonstrating a striking loss of diurnal BP rhythmicity in homozygous null rats that was restored by reducing dietary salt intake.

It has been demonstrated in the mouse SAME model (14) that heterozygous Hsd2^+/-^ mice show a blunted response to salt load handling, going transiently into positive salt balance and exhibiting salt sensitive hypertension (15). Unlike wild-type mice, ENaC activity persists on high salt diet (16) despite an appropriate reduction in aldosterone. Homozygous knockout mice initially show an increased epithelial sodium channel (ENaC) activity, associated with impaired sodium excretion (13) but by 3 months of age, ENaC trafficking to the apical membrane is abolished, and the mice become salt wasting (aldosterone escape). Interestingly, both this and the hypertension are prevented by glucocorticoid receptor (GR) antagonism, indicating that it is GR, rather than MR, which affects the sodium mishandling in the mouse model. Older Hsd2^-/-^ mice develop nephrogenic diabetes insipidus (a defect in urine concentrating ability(25)) and a progressive loss of functional renal medulla (14). Here we show that the rat model of SAME (19) shows significant differences from the mouse.

A detailed telemetric analysis failed to distinguish between Hsd2^+/-^ and F344 control rats. In the present study, heterozygous rats were found not to be significantly hypertensive compared to F344 controls. When WT and Hsd2^+/-^ rats were put on a 3% Na diet they increased MABP equally by ∼6mmHg, suggesting that, unlike the SAME mouse model, they are equivalently susceptible to salt-induced hypertension, at least on the F344IcoCrl background. This suggests that the F344 rat handles salt load in a different way to the mouse. Basset et al (26) previously reported that F344 rats have reduced dipping compared to other rat strains such as Wistar. This was confirmed in the present study (Table 1), where WT animals showed only ∼3.5mmHg LD difference on both the 0.3% and 3% Na diets and an LD phase ratio of 3% (rather than the 10% attributed to other rat strains).

Homozygous rats were found to be severely hypertensive on the 0.3% Na diet (Table 2; Fig2a – treatment 1). Notably, the homozygotes had virtually no dipping of MABP during the inactive daytime period, compared to controls on the same diet. In fact, Hsd2^-/-^ animals regularly registered a higher average MABP during the light than the dark period. This did not reflect heart rate (Fig. 2b – treatment 1) or locomotor activity, which both exhibited circadian rhythm as expected. However, the heart rate was significantly slower than controls.

When the Hsd2^-/-^ rats were transferred to a 0.03% Na diet, their blood pressure dropped by 40 mmHg and a robust circadian rhythm of light-dark MABP developed (Fig2a – treatment 2). The LD phase ratio increased to levels higher than those observed in the WT animals on either 0.3% or 3% salt (Table 2). Non-dipping in healthy human individuals has been shown to be associated with elevated myocardial re-polarisation lability and impaired baroreflex function, both of which are suggestive of autonomic nervous system dysfunction (27). It has previously been observed in some human studies that circadian variation in blood pressure is affected by diet (28) and a reduction in sodium intake can rectify the non-dipper phenotype (9,29). It is possible that the F344IcoCrl rat strain is already primed for non-dipping, given the low LD phase ratio observed in control animals on normal (0.3% Na) rat chow, and its observed sensitivity to salt. The extreme salt sensitive hypertension of the Hsd2^-/-^ rats may relate to exacerbated ANS dysfunction, causing non-dipping even on 0.3% Na diet. Reduction of Na to 0.03% may dramatically reduce the blood pressure by alleviating the ANS dysfunction.

Treatment of Hsd2^-/-^ rats with the non-selective inverse agonist of the α1-adrenoreceptor, prazosin, which blocks noradrenalin-induced vasoconstriction of vascular smooth muscle cells, caused a further drop in blood pressure of ∼20mmHg, on 0.03% Na diet but the exaggerated dipping phenotype was no longer maintained. This indicates that the sympathetic nervous system contributes to the hypertensive phenotype and suggests that the non-dipping phenotype results from a complex interaction between the sympathetic and parasympathetic nervous systems.

Recently, Sueta et al (30) reported that the SHRcp strain of rat, a sub-strain of SHR, which carries a spontaneous mutation in the leptin receptor gene, exhibits metabolic syndrome and a non-dipper type of hypertension. The high levels of urinary norepinephrine and aldosterone in this model suggested enhanced sympathetic tone and impaired baroreflex function. Treatment with azilsartan, an AT1 receptor antagonist, reduced blood pressure, urinary norepinephrine and urinary aldosterone in a dose-dependent manner, suggesting that the hypertension was partly caused by angiotensin II-mediated autonomic dysfunction in this model. However, azilsartan had little effect on the non-dipper phenotype (see Fig. 5 in reference (30)). The evident parallels with our Hsd2 knockout model, which has ‘apparent’ mineralocorticoid excess and a non-dipper phenotype, are clear. However, in the present study, hypertension and non-dipping were improved and rectified, respectively, through a simple reduction in dietary salt.

Despite the effect of salt treatment on MABP in the F344 controls and Hsd2^+/-^ rats, an increase in sodium did not effect a change in heart rate. The Hsd2^-/-^ animals, however, exhibited a much slower heart rate on the 0.3% diet, which increased with decreasing sodium. Interestingly, prazosin caused a further increase in heart rate to wild type levels. Recently, it was demonstrated that high salt with saline causes a decrease in heart rate in Sprague Dawley rats, but no change in heart rate in Dahl salt-sensitive rats. The latter were, however, shown to have increased renal sympathetic nerve activity on high salt treatment (31). In the present study, reduction in salt led to an increase in heart rate in the Hsd2^-/-^ rats. This indicates that heart rate also results from a complex interaction between the sympathetic and parasympathetic nervous systems, which varies between strains of rat.

It has previously been shown, using the recurrence plot, that non-linear indexes of blood pressure and heart rate variability respectively reflect sympathetic and parasympathetic involvement in normotensive (32) and hypertensive (SHR) rats (33). Though the relation between heart rate, heart rate variability and vagal nerve stimulation is not clear, rat studies using various periodic or stochastic stimulatory protocols on the right vagal nerve (34) have demonstrated clear effects on heart rate dynamics.

In the present study, using Poincare analysis, we demonstrated an enhancement of the circadian variation in SD1 on reduction of salt in the diet, suggesting an association between parasympathetic nervous system dysfunction and the observed non-dipper phenotype. Carvalho et al (35) presented compelling evidence that the non-dipper phenotype may result, at least in part, from a decrease in parasympathetic function. They demonstrated that patients with impaired parasympathetic ANS, but intact sympathetic ANS had an attenuated reduction in nocturnal BP. Dysfunction of both parasympathetic and sympathetic ANS led to a complete loss of circadian variation. Recently, circadian changes in various HRV parameters were demonstrated in healthy control subjects, which were depressed in patients with drug-resistant epilepsy (36).

Hsd11b2 is expressed in the placenta (37), where it plays a critical role in preventing exposure of the fetus to high circulating levels of glucocorticoid (38). Apart from the effect of this on salt handling in the skin of Hsd2^-/-^ knockout rats (39), it is possible that exposure to aberrant Na^+^ levels in the developing neonate leads to abnormal ANS function. Of the many locations in the rat brain where sodium appetite is thought to be regulated, the nucleus of the solitary tract uniquely contains a subset of aldosterone-sensitive neurons that co-express Hsd11b2 (40) and respond specifically to sodium deprivation. Apart from vagal innervation from the abdominal viscera (41) to the HSD2 neurons, there is no evidence for direct afferent or efferent connections to or from these neurons and sympathetic or parasympathetic centers (42) but rather to sites in the fore-brain implicated in sodium appetite. The intriguing involvement of salt, blood pressure, sympathetic and parasympathetic dysfunction in the SAME rat model, strongly indicate a complex interaction between diet, blood pressure and autonomic nervous system. Detailed investigation should allow us to understand these interactions, which will be directly applicable to the human condition.

## Acknowledgements

We wish to acknowledge the technical assistance of Mrs. G. Brooker and Dr Jon Manning and helpful discussions with Dr Christopher Kenyon and Dr G. Culshaw.

## Sources of Funding

Authors are grateful for funding from the British Heart Foundation Centre for Research Excellence (RE/08/001/23904).

## Conflict(s) of Interest/Disclosure(s) Statement

None

## Notes

### Competing Interest Statement

The authors have declared no competing interest.

